# Integrated Single Cell Atlas of Endothelial Cells of the Human Lung

**DOI:** 10.1101/2020.10.21.347914

**Authors:** Jonas C. Schupp, Taylor S. Adams, Carlos Cosme, Micha Sam Brickman Raredon, Norihito Omote, Sergio Poli De Frias, Kadi-Ann Rose, Edward Manning, Maor Sauler, Giuseppe DeIuliis, Farida Ahangari, Nir Neumark, Yifan Yuan, Arun C. Habermann, Austin J. Gutierrez, Linh T. Bui, Kerstin B. Meyer, Martijn C. Nawijn, Sarah A. Teichmann, Nicholas E. Banovich, Jonathan A. Kropski, Laura E. Niklason, Dana Pe’er, Xiting Yan, Robert Homer, Ivan O. Rosas, Naftali Kaminski

**Affiliations:** Pulmonary, Critical Care and Sleep Medicine, Yale School of Medicine, New Haven, CT, USA; Department of Biomedical Engineering, Yale University, New Haven, CT, USA; Vascular Biology and Therapeutics, Yale University, New Haven, CT, USA; Department of Medicine, Baylor College of Medicine, Houston, TX, USA; Division of Internal Medicine, Mount Sinai Medical Center, Miami Beach, FL, USA; VA Connecticut Healthcare System, West Haven, CT, USA; Department of Anesthesiology, Yale University, New Haven, CT, USA; Division of Allergy, Pulmonary and Critical Care Medicine, Department of Medicine, Vanderbilt University Medical Center, Nashville, TN, USA; Translational Genomics Research Institute, Phoenix, AZ, USA; Cellular Genetics, Wellcome Sanger Institute, Hinxton, Cambridge, UK; Department of Pathology and Medical Biology, University Medical Center Groningen, University of Groningen, Groningen, the Netherlands; Groningen Research Institute for Asthma and COPD, University Medical Center Groningen, University of Groningen, Groningen, the Netherlands; Theory of Condensed Matter Group, Cavendish Laboratory/Department of Physics University of Cambridge, Cambridge, UK; Department of Veterans Affairs Medical Center, Nashville, TN, USA; Department of Cell and Developmental Biology, Vanderbilt University, Nashville, TN, USA; Program for Computational and Systems Biology, Sloan Kettering Institute, Memorial Sloan Kettering Cancer Center, New York, NY, USA; Department of Pathology, Yale University School of Medicine, New Haven, CT, USA; Pathology and Laboratory Medicine Service, VA Connecticut HealthCare System, West Haven, CT, USA

## Abstract

**Background:** Despite its importance in health and disease, the cellular diversity of the lung endothelium has not been systematically characterized in humans. Here we provide a reference atlas of human lung endothelial cells (ECs), to facilitate a better understanding of the phenotypic diversity and composition of cells comprising the lung endothelium, both in health and disease.

**Methods:** We reprocessed control single cell RNA sequencing (scRNAseq) data from five datasets of whole lungs that were used for the analysis of pan-endothelial markers, we later included a sixth dataset of sorted control EC for the vascular subpopulation analysis. EC populations were characterized through iterative clustering with subsequent differential expression analysis. Marker genes were validated by immunohistochemistry and *in situ* hybridization. Signaling network between different lung cell types was studied using connectomic analysis. For cross species analysis we applied the same methods to scRNAseq data obtained from mouse lungs.

**Results:** The six lung scRNAseq datasets were reanalyzed and annotated to identify over 15,000 vascular EC cells from 73 individuals. Differential expression analysis of EC revealed signatures corresponding to endothelial lineage, including pan-endothelial, pan-vascular and subpopulation-specific marker gene sets. Beyond the broad cellular categories of lymphatic, capillary, arterial and venous ECs we found previously indistinguishable subpopulations; among venous EC we identified two previously indistinguishable populations, pulmonary-venous ECs (COL15A1^neg^) localized to the lung parenchyma and systemic-venous ECs (COL15A1^pos^) localized to the airways and the visceral pleura; among capillary EC we confirmed their subclassification into recently discovered aerocytes characterized by EDNRB, SOSTDC1 and TBX2 and general capillary EC. We confirmed that all six endothelial cell types, including the systemic-venous EC and aerocytes are present in mice and identified endothelial marker genes conserved in humans and mice. Ligand-Receptor connectome analysis revealed important homeostatic crosstalk of EC with other lung resident cell types. Our manuscript is accompanied by an online data mining tool (www.LungEndothelialCellAtlas.com).

**Conclusion:** Our integrated analysis provides the comprehensive and well-crafted reference atlas of lung endothelial cells in the normal lung and confirms and describes in detail previously unrecognized endothelial populations across a large number of humans and mice.

## Introduction

The most critical function of the circulatory system is to provide tissues throughout the body with a constant supply of oxygenated blood. In mammals, the circulatory system accomplishes this feat by performing gas exchange with the environment exclusively in the lung. Consistent with its hyperspecialized role, the lung is a highly vascularized organ uniquely comprised of two circulations ^1^. Systemic circulation is primarily restricted to the large airways and the pleura, to which it supplies oxygenated blood. In contrast, the pulmonary circulation supplies oxygen-depleted blood to the lung parenchyma to perform gas exchange. The contrasting physiological functions of these circulations, as well as the distinct cellular niches they each reside in underscores the diverse and unique roles played by endothelial cells in the lung.

It had been thought that the endothelium mainly served a static role as a physical barrier between the blood, the air and the stromal tissue^2^. However, this ‘static’ characterization of the endothelium has more recently been replaced with a more ‘dynamic’ understanding of the cell type due to its rich functional heterogeneity and advancements in our understanding of endothelial cell physiology^3, 4^. The endothelium is metabolically active and has been found to play a key role in processes governing inflammation, leukocyte trafficking, gas and nutrient exchange, hemostasis, angiogenesis, vasoconstriction/vasodilation and endocrine signaling^4, 5^. In addition, lung endothelium experiences constant stretching while breathing and is persistently exposed to substances of the external environment. Dysfunction of the lung endothelium has been described in numerous clinical conditions including acute respiratory distress syndrome (ARDS), chronic obstructive pulmonary disease (COPD), pulmonary fibrosis, pneumonia, autoimmune conditions, pulmonary hypertension and others^2^.

Despite the obvious importance of endothelial cells to lung function in health and disease, a detailed reference atlas of endothelial cells in the healthy human lung is missing. We created an integrated single-cell atlas of human lungs to identify endothelial cell populations, characterize them by prototypic gene expression profiles and explore their role in tissue homeostasis. To identify cross species conserved cell populations and markers, as well as key differences between the mouse and the human, we generated an integrated murine endothelial atlas and compared human lung endothelial cell (EC) populations to that of mice. Our manuscript is accompanied by a publicly accessible online tool to interactively explore the data (www.LungEndothelialCellAtlas.com).

## Methods

Detailed methods can be found in the supplement.

## Results

### Integrated single cell atlas of the human lung

In order to profile endothelial cell populations in the human lung, we collected, reprocessed and integrated single cell RNA sequencing (scRNAseq) data of five publicly available datasets through a common computational pipeline (details see Supplemental Methods): Vanderbilt-TGen^6^ (n=10), WSI-Groningen^7^ (n=10), Northwestern^8^ (n=8), Leuven VIB^9^ (n=6) and Yale-Baylor^10^ (n=34, including samples from four previously unpublished additional subjects, Supp. Table S1). The median age of subjects was 50 years (interquartile range (IQR) 32 - 61 years), and 29 out of 68 were female (for basic characteristics by cohort, see Supp. Table S2). In all cohorts, single cell barcoding had been performed using the 10x Single Cell RNA 3prime sequencing kits based on v2 chemistry with the notable exception of the Vanderbilt-TGen cohort which had utilized the 5prime technology. Four samples were derived from airway biopsies, and all others from dissociated distal lung samples. Raw scRNAseq data from all 68 human lung samples were aggregated and collectively processed, resulting in the lung dataset consisting of 278,648 cells including 12,563 vascular and lymphatics ECs from 66 of the 68 subjects. We identified 37 cell types that aligned with canonical marker gene expression signatures of established cell types (see figure 1B, 2A, Supp. Figure S1A and Supp. Table S3). Importantly, reported marker genes were universally detected across cell types, all scRNAseq assay schemes, all cohorts and most subjects, and therefore do not represent subject-specific gene expression. Technical summaries of sample preprocessing results can be found in Supp. Table S4.

**Figure 1.**
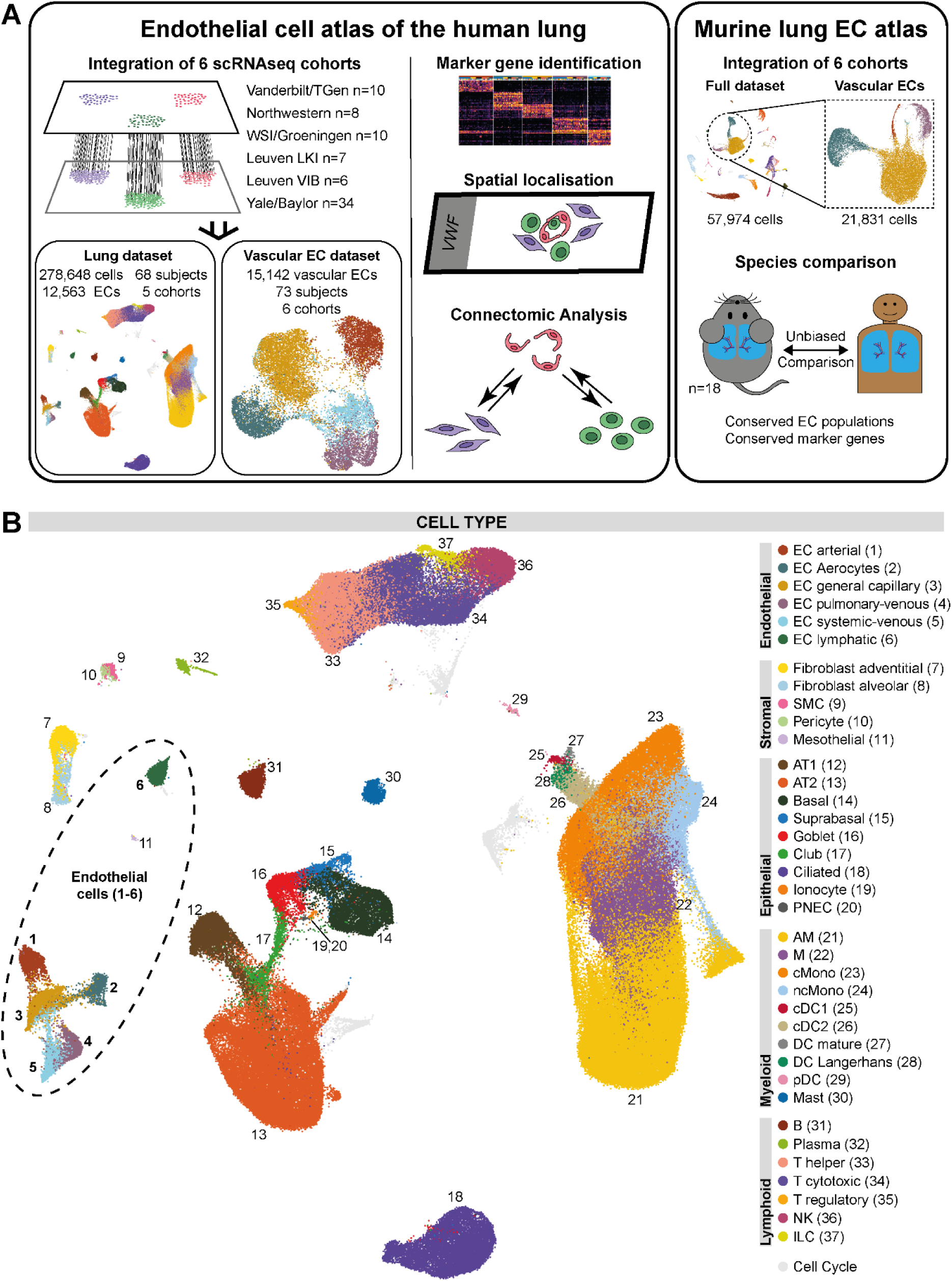
Overview of the study design and the integrated dataset. **(A)** Overview of study design. (i) Integration of scRNAseq datasets including four previously unreported samples. (ii) Annotation of cell types. (iii) Identification of markers genes at several levels of lineage. (iv) Spatial localization by immunohistochemistry and in situ hybridization. (v) Connectomic analysis of cell-cell-communication. (vi) Identification of conserved EC populations and marker genes in human and mice. **(B)** Uniform Manifold Approximation and Projection (UMAP) representation of 278,648 cells from 68 control lungs; each dot represents a single cell, and cells are labeled as one of 37 discrete cell types. For UMAPs colored by cohort and subject, please refer to Supp. Figure S1A. AM: alveolar macrophage; AT1/2: alveolar cell type 1/2; B: B cell; cDC1/2: classical dendritic cell type 1/2; pDC: plasmacytoid dendritic cell; DC: dendritic cell; EC: endothelial cell; M: macrophage; NK: natural killer; ILC. innate lymphoid cell; PNEC: pulmonary neuroendocrine cell; scRNAseq: single cell RNA sequencing; SMC: smooth muscle cell; T: T cell.

### Lung endothelial cell markers, assessed at levels of lineage, anatomy, and cell-type

We identified 12,563 ECs within the integrated lung dataset based on the canonical endothelial markers PECAM1 (CD31), CDH5 (VE-cadherin), CLDN5, and ERG (Figure 2A,2B). We identified 147 pan-endothelial marker genes, defined as genes significantly expressed in all vascular and lymphatic EC populations compared to all other cell types (Supp. Table S3). Pan-endothelial markers include genes coding for proteins associated with endothelial-intrinsic structures like the glycocalyx (PODXL, ST6GALNAC3, GALNT18), caveolae (CAV1, CAV2, CAVIN1, CAVIN2), focal adhesion structures (ITGA5, ITGB1, LAMB2, AKT3, PXN, MYL12A, MYL12B, PRK2) and adherens junctions (AFDN, TJP1, PTPRM) (Figure 2A, Supp. Table S3). All EC subpopulations express common transcription factors (ERG, ETS1, FLI1, SOX18) and regulators of endothelial migration (ANXA3, SASH1, LDB2, PTK2, MEF2C, STARD13). Other pan-endothelial markers include angiopoietin receptor TIE1, subunits of the receptors for calcitonin-gene-related peptide and adrenomedullin CALCRL and RAMP2/RAMP3, as well as general endothelial receptors ROBO4 and S1PR1 (Figure 2A, Supp. Table S3).

**Figure 2.**
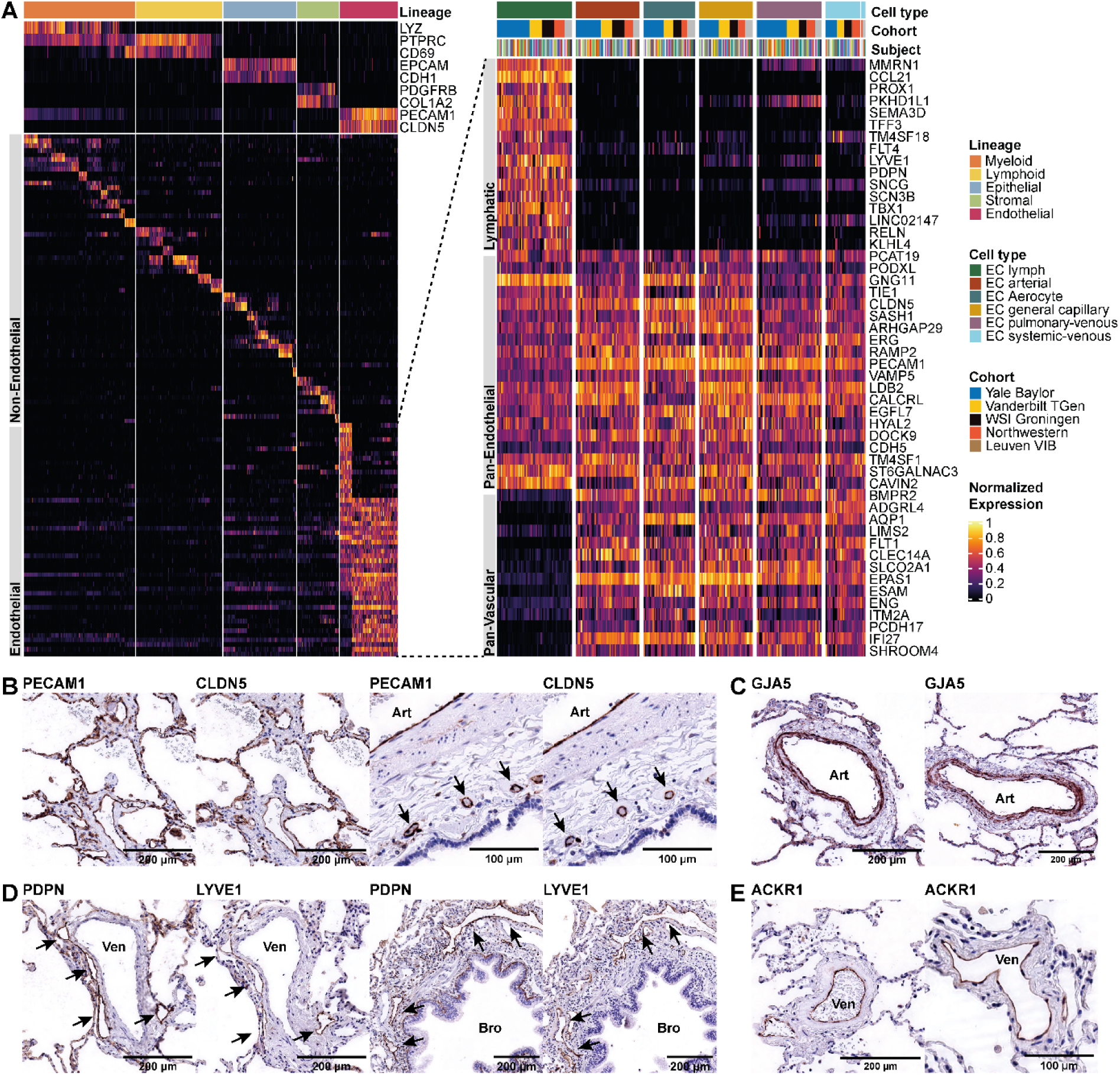
Assessing coarse-granular endothelial heterogeneity of the human lung by scRNAseq. **(A)** Heat map of marker genes for all 29 non-EC cell types and of lymphatic cells, as well as pan-endothelial (specifically expressed in all six EC populations) and pan-vascular (specifically expressed in all EC populations but lymphatic ECs) marker genes. Each column represents the average expression value for one subject, grouped by cell type and cohort. All gene expression values are unity normalized from 0 to 1 across rows. For an enlarged heatmap of non-EC cell types, please refer to Supp. Figure S1B. **(B)** IHC staining of pan-endothelial markers CLDN5 and PECAM1, with positive brown staining in capillaries and veins in the first figure pair and arterial (“Art”) and peribronchial vessels (arrows) in the second figure pair. **(C)** IHC staining of arterial marker GJA5 with positive staining of an arteriole (“Art”) in brown. **(D)** IHC staining of lymphatic markers PDPN and LYVE1, with positive brown staining in lymphatic vessels (arrows) surrounding a larger vein (“Ven”) in the first figure pairs and a bronchus (“Bro”) in the second figure pair. As predicted by the transcriptomic scRNAseq data, PDPN also stains basal cells and AT1. **(E)** IHC staining of venous marker ACKR1 with positive staining of venules (“Ven”) in brown. AM: alveolar macrophage; AT1/2: alveolar cell type 1/2; B: B cell; cDC1/2: classical dendritic cell type 1/2; pDC: plasmacytoid dendritic cell; DC: dendritic cell; EC: endothelial cell; M: macrophage; NK: natural killer; ILC: innate lymphoid cell; PNEC: pulmonary neuroendocrine cell; SMC: smooth muscle cell; T: T cell.

Lung ECs separated into two broad populations in UMAP space (Figure 1A, 2A, 2D, Supp. Table S3): vascular ECs (10,469 cells from 66 subjects) and lymphatic ECs (2,094 cells from 64 subjects). Lymphatic vessels, lined by lymphatic ECs, absorb interstitial fluid, and carry it as lymph to a lymph node or collecting vein. Lymphatic ECs were identified based on the expression of canonical lymphatic markers PROX1^11^, LYVE1^11^, FLT4^11, 12^ PDPN^12, 13^ (Figure 2A, Supp. Table S3). Immunohistochemistry (IHC) staining for lymphatic markers LYVE1 and PDPN revealed lymphatic vessels in the distal lung (lymphatic capillaries) as well as surrounding large vascular vessels and bronchi (Figure 2D). Lymphatic ECs express secreted proteins like chemokine CCL21^12^ mediating homing of T-cells and semaphorins SEMA3A and SEMA3D^14^, both necessary for lymphatic vessel maturation. Furthermore, lymphatic marker genes comprise transcription factors TBX1, HOXD3, NR2F1, NR2F2 and receptors KDR, GPR182, TEK.

In contrast, blood vessels are lined by vascular ECs and carry blood from and to the heart. Pan-vascular marker (n=142) expressed across the remaining, non-lymphatic endothelial cells include transmembrane glycoproteins such as ENG, PCDH17, CLEC14A, CLEC1A, ESAM, ITM2A, the cell surface receptors BMPR2, FLT1, ADGRL4, VIPR1, PLXNA2, FZD4, IL4R, IL15RA, transporter and channel proteins such as SLCO2A1, SLCO4A1, AQP1 as well as the transcription factors EPAS1, GATA2, FOXF1, ETS2 (Figure 2A, Supp. Table S3).

### Identification of arterial, venous, and capillary endothelial cells

To focus on vascular EC, we generated a non-lymphatic EC dataset. To increase the power of our analysis, we included control vascular ECs from a sixth scRNAseq dataset^15^ (Leuven LKI, n=7 subjects) of sorted, CD45^neg^/CD31^pos^ lung ECs that brought the total number of vascular ECs to 15,142 from 73 subjects. The cohort “Leuven LKI” had not been included in the initial lung dataset to avoid batch effects due to missing or selectively sorted non-ECs within this cohort.

We identified three broad vascular EC subtypes: arterial (n=2,610 cells), capillary (n=7,920 cells) and venous (n=4,612) ECs based on expression of canonical marker genes (Figure 3A, 3B, Supp. Table S5). Arterial ECs were recognized by EFNB2^16-19^, SOX17^17, 20^, BMX^16, 21^, SEMA3G^16^, HEY1^17, 18^, LTBP4^22^, FBLN5^22^, GJA5^17^ and GJA4^17^. Capillary ECs were identified by canonical markers CA4^12, 23^ and PRX^24^, as well as organism-wide capillary markers RGCC^12^, SPARC^12^ and SGK1^12^. Venous EC were identified through the canonical transcription factor NR2F2^17-19^ (COUP-TFII), as well as VCAM1^12, 19^, ACKR1^25^ (Figure 2E) and SELP^25^ expression. Capillary and venous ECs separated further into two distinct populations each - venous ECs into pulmonary-venous and systemic-venous ECs and capillary ECs into aerocytes and general capillary ECs, respectively – for which a detailed characterization will be given in the two subsequent paragraphs (for makeup of vascular ECs, see Supp. Fig. 2A).

**Figure 3.**
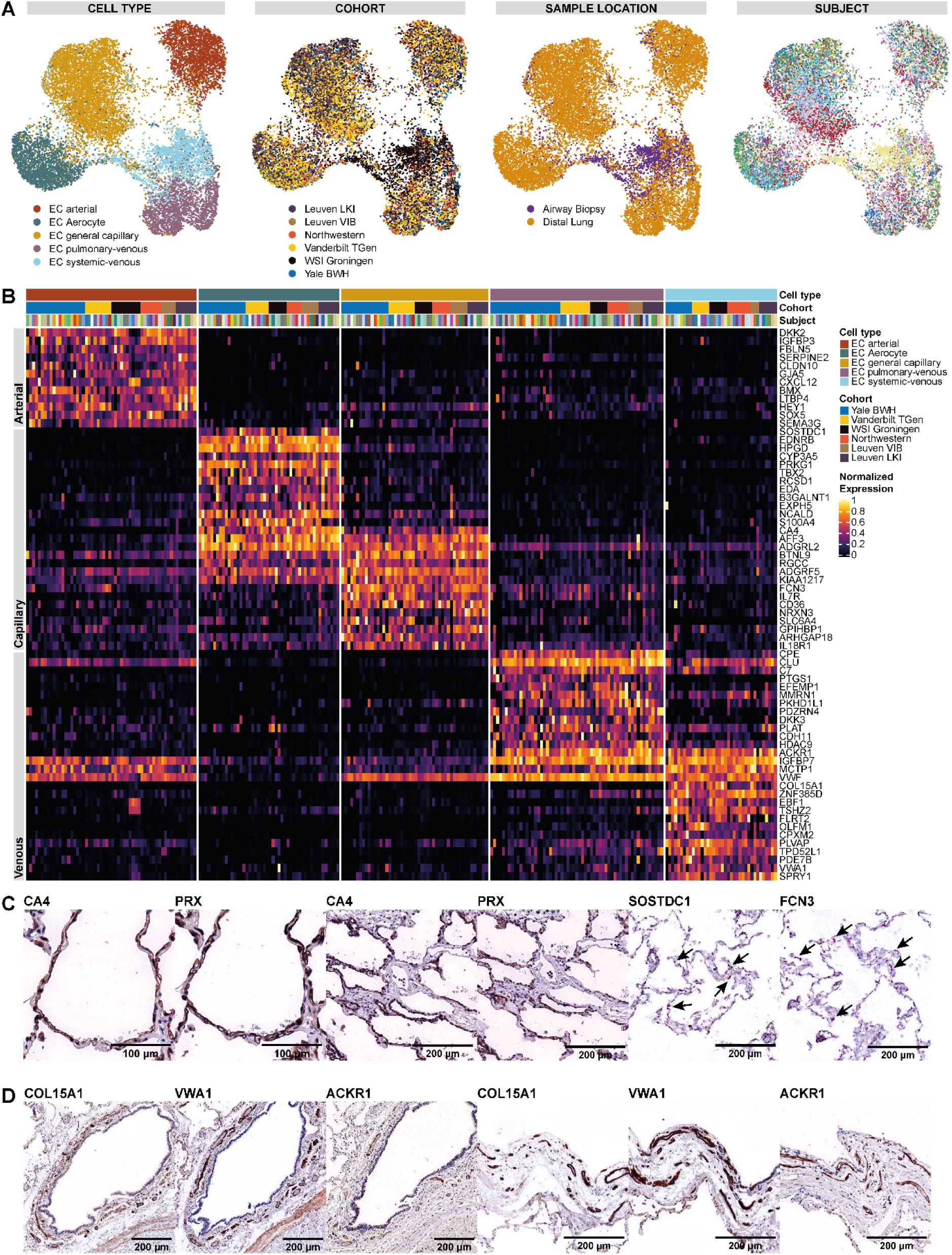
Assessing fine-granular endothelial heterogeneity of the human lung by scRNAseq. **(A)** UMAPs of 15,142 endothelial cells from 73 control lungs. Each dot represents a single cell, and cells are labelled – from left to right - by (i) cell type, (ii) cohort, (iii) sample location and (iv) subject. In the UMAP colored by subjects, each color represents a distinct subject. **(B)** Heat map of marker genes of all five EC populations. Each column represents the average expression value for one subject, grouped by cell type and cohort. All gene expression values are unity normalized from 0 to 1 across rows. **(C)** IHC staining of capillary markers PRX and CA4 with positive brown staining in capillaries, in higher magnification in the first two figure pairs focusing on an alveolus and a lower magnification in the second two figure pairs in which the negative staining of a larger central vessel can be appreciated. The last figure pairs are in situ RNA hybridization stains of markers specific to the capillary subpopulations with SOSTDC1 staining aerocytes ECs (arrows) and FCN3 staining general capillary ECs (arrows) with positive staining in red. **(D)** IHC staining of systemic-venous EC markers COL15A1, VWA1 and ACKR1 with positive brown staining in small vessels surrounding a larger bronchus in the first three subfigures and in small vessels of the visceral pleura in the last three subfigures.

Beyond canonical marker concordance, vascular EC subpopulations could be identified based on their distinct gene signatures associated with their respective cellular roles. Arterial ECs lining pulsating arteries and arterioles are exposed to relevant transmural pressure and shear-stress. To withstand these forces, arterial ECs express genes encoding for tight and gap junction proteins such as CLDN10, GJA5 (Figure 2C), GJA4, FBLIM1, extracellular matrix proteins that contribute to wall elasticity and strength such as FBLN5, FBLN2, MGP, BGN, LTBP4, LTBP1, FN1 and the protease inhibitors SERPINE2^26^ and CPAMD8 (Figure 3B, Supp. Table S5). Arterial ECs are active secretory cells and express signaling molecules such as CXCL12, EFBN2, SEMA3G^27^, VEGFA, TAFA2 and enzymes of the nitric oxide pathway involved in modulating vascular tonus like NOS1, PDE3A and PDE4D. Maintenance of their arterial identity is accomplished through the expression of Wnt signaling modulator DKK2, the Notch ligand DLL4, and the transcription factors HEY1, SOX5, SOX17, HES4 and PRDM16 (Figure 3B, Supp. Table S5).

The capillary network surrounds millions of alveoli in the lungs to enable gas exchange. Capillary ECs express genes coding for enzymes related to gas exchange including CA4^12, 23^, CYB5A, multiple receptors including ACVRL1/TMEM100, ADGRF5, ADGRL2, F2RL3, IFNGR1, VIPR1, ADRB1, intracellular signaling molecules like ARHGAP6, IFI27, PREX1, PRKCE, SGK1^12^, SH2D3C, SORBS1, structural proteins PRX^24^, SPARC^12^, EMP2, ITGA1, SLC9A3R2, transcription regulators AFF3 and MEIS1, and exhibit the highest MHC class I protein expression of all endothelial cells (Figure 3B, Supp. Table S5).

Venules and veins return blood to the heart. In contrast to arteries, veins are thin-walled and subject to low shear forces which makes in particular the postcapillary venules the primary location for leukocyte extravasation. To further facilitate this, venous ECs express genes encoding for proteins associated with diapedesis of leukocytes like VCAM1^12, 19^, SELP^25^, SELE^25^ (Figure 3B, Supp. Table S5) as well as ACKR1^25^ for unspecific transcytosis of chemokines. Venous EC are further characterized by the major transcription factor NR2F2^17-19^, as well as genes coding for secreted (ADAMTS9, IGFBP7), nuclear (HDAC9, RORA) and membrane proteins (ACTN1, LDLRAD3, LDLRAD4, LRRC1) (Supp. Table S5).

### scRNAseq characterization of two previously indistinguishable capillary endothelial cell types

scRNAseq reveals that the previously and consistently observed patchy or mosaic histological staining patterns of VWF, THBD and EMCN^15, 28-32^ as well as EDN1 and EDNRB1^33^ in lung capillaries can be attributed to two discernable capillary populations: a VWF^neg^/EMCN^high^/EDNRB1^pos^ population and a VWF^pos^/EMCN^low^/EDN1^pos^ population (Figure 3A,B). Following nomenclature proposed by Gillich *et al*.^34^ we refer to these two microvasculature ECs as aerocytes and general capillary ECs.

Aerocytes (n=2,317) can be distinguished by their expression of genes encoding for the Endothelin Receptor EDNRB, the transcription factors TBX2, FOXP2, the pattern recognition receptors CLEC4E, SPON2, and signaling mediators such as PRKG, CHRM2, S100A4, EDA (Figure 3B, Supp. Table S5). Aerocytes are the only EC type expressing prostaglandin-degrading HPGD, which is responsible for the first-pass deactivation of most prostaglandins as they pass through the vascular bed of the lung^35^. Aerocytes specifically express SOSTDC1, antagonist to BMPR2, which itself is expressed in all ECs and its mutations are associate with pulmonary arterial hypertension. In contrast to all other ECs, aerocytes do not express major components of endothelial-specific Weibel-Palade bodies (VWF, SELP, EDN), storage granules essential for hemostasis and extravasation of leukocytes. Additionally, aerocytes are characterized by the absence of expression of genes otherwise expressed in all other ECs including THBD, CD93, PTPRB, ANO2, CRIM1, MECOM, LIFR, PALMD, GNA14, CYYR1 and LEPR. Aerocytes are rarer in the human lung compared to general capillary ECs and were profiled in the median per subject in a ratio of aerocytes to general capillary ECs of 0.48:1 (interquartile range (IQR) 0.26 – 1.16, Supp. Figure 2B).

General capillary ECs (n= 5,603) are the most generic vascular cell type, as they specifically express only 140 genes, compared to arterial (n=373) or venous (n=280) EC or aerocytes (n=203). Nevertheless, the general capillary EC population can be identified by the expression of genes related to transcytosis of LDL and other lipids such as GPIHBP1^12^ and CD36^31^, innate immune response including FCN3, BTNL9, BTNL8, CD14, and cytokine receptors such as IL7R, IL18R1 (Figure 3B, Supp. Table S5). We localized both capillary EC types by immunohistochemistry using CA4 and PRX as common capillary markers and by in-situ hybridization using the subtype specific markers SOSTDC1 for aerocytes and FCN3 for general capillary ECs (Figure 3C). In the control lungs, both consistently localize to and intermingle in the endothelium of the alveolar wall.

### Systemically perfused vessels of bronchi and the visceral pleura are lined by a COL15A1-positive endothelial cell type

Venous ECs divide into a COL15A1^pos^ and a COL15A1^neg^ subpopulations. Immunohistochemical analysis using the markers COL15A1 and VWA1 revealed that COL15A1^pos^ ECs localized to systemically supplied vessels of the bronchial vascular plexus – as we previously reported^10^ - and of the visceral pleura (Figure 3D), while vessels supplied by the pulmonary circulation remained negative. In contrast, the pan-venous marker ACKR1 stained both pulmonary and systemic venules (Figure 2E, 3D). Furthermore, four scRNAseq samples from biopsies of the systemically supplied large airways contributed almost exclusively cells to the COL15A1^pos^ population (Figure 3A). Based on the IHC staining and the tissue source of the samples, we therefore were able to assign two venous subpopulations to the systemic and pulmonary circulation and thus call them systemic-venous ECs (COL15A1^pos^, n=2,291) and pulmonary-venous ECs (COL15A1^neg^, n= 2,321). The integrated dataset allowed us to identify additional genes specific to systemic-venous ECs in addition to COL15A1, which is a multiplexin collagen linking basement membrane and its underlying connective tissue. We also identified the extra-cellular matrix protein VWA1 as well as PLVAP, that encodes an endothelial cell-specific protein that forms the stomatal and fenestral diaphragms of blood vessels and regulates basal permeability, leukocyte migration and angiogenesis (Figure 3B, Supp. Table S5). Systemic-venous ECs express genes that encode for specific sets of transcription factors including EBF1, EBF3, MEOX1, MEOX2, ZNF385D, the cell surface receptors TACR1, ROBO1, CYSLTR1, the peptidases CPXM2 and MMP16, the phosphodiesterases PDE7B, PDE2A, as well as antagonist of fibroblast growth factor signaling SPRY1.

While aerocytes are characterized by their ability to break down prostaglandins (HPGD) and their lack of hemostatic Weibel-Palade bodies, pulmonary-venous EC - which are located downstream - express genes encoding for proteins associated with prostaglandin synthesis including PTGS1 and PTGIS, and with complement and coagulation cascades such C7, PLAT, PROCR, THBD, C1R, CLU (Figure 3B, Supp. Table S5). Pulmonary-venous ECs express the most specific pulmonary-venous gene CPE as well as Wnt modulator DKK3, which contrasts to the DKK2 expression in arterial ECs. Genes coding for structural proteins specific to pulmonary-venous ECs include EFEMP1 and CDH11. Overlapping with lymphatic ECs, but expressed at lower levels compared to them, pulmonary-venous ECs express factor V/Va-binding and extracellular matrix protein MMRN1^12^ and PKHD1L1 (Figure 3B, Supp. Table S5).

### Endothelial Cell subpopulations’ connectivity with other lung cell populations

Complex tissues like the lung have distinct cellular niches that are tightly regulated by intercellular communications. To study the role of ECs in tissue homeostasis within the human lung, we performed a connectomic analysis that mapped the averaged gene expression per cell type to known ligand-receptor interactions catalogued in the NicheNet database^36, 37^. We focused our analysis on potential communication pathways between EC and non-EC cell types. Due to the complexity of the cell-cell signaling connectome, we can only present a limited fraction of our findings here (full connectome, see Supp. Table S6). All ligand-receptor interaction networks can be explored on www.LungEndothelialCellAtlas.com.

We first identified the intercellular signaling patterns wherein ECs are “senders” of ligands (Figure 4A). Arterial ECs are the most active EC subpopulation in this respect, expressing Notch ligands (DLL4, JAG1 and JAG2) which can be sensed by NOTCH3 in pericytes and smooth muscle cells (SMCs) as a part of a signaling axis that is crucial for arterial EC specification, as well as proliferation and differentiation of supporting pericytes and vascular SMCs^38^. Arterial ECs, and to a lesser extent venous and general capillary ECs express EDN1 which can be sensed by pericytes, SMC and alveolar fibroblasts through EDNRA; this is a vasoconstrictive axis^39^ implicated in pulmonary hypertension. Arterial endothelial cells also secrete CXCL12, a chemokine that facilitates homing, migration, and survival of immune cells through binding to CXCR4 which is expressed by most lymphoid (T regulatory, T helper, T cytotoxic, B cells, and innate lymphoid cells) and dendritic (plasmacytoid (pDC), mature DC, and Langerhans DC) cells^40^.

**Figure 4.**
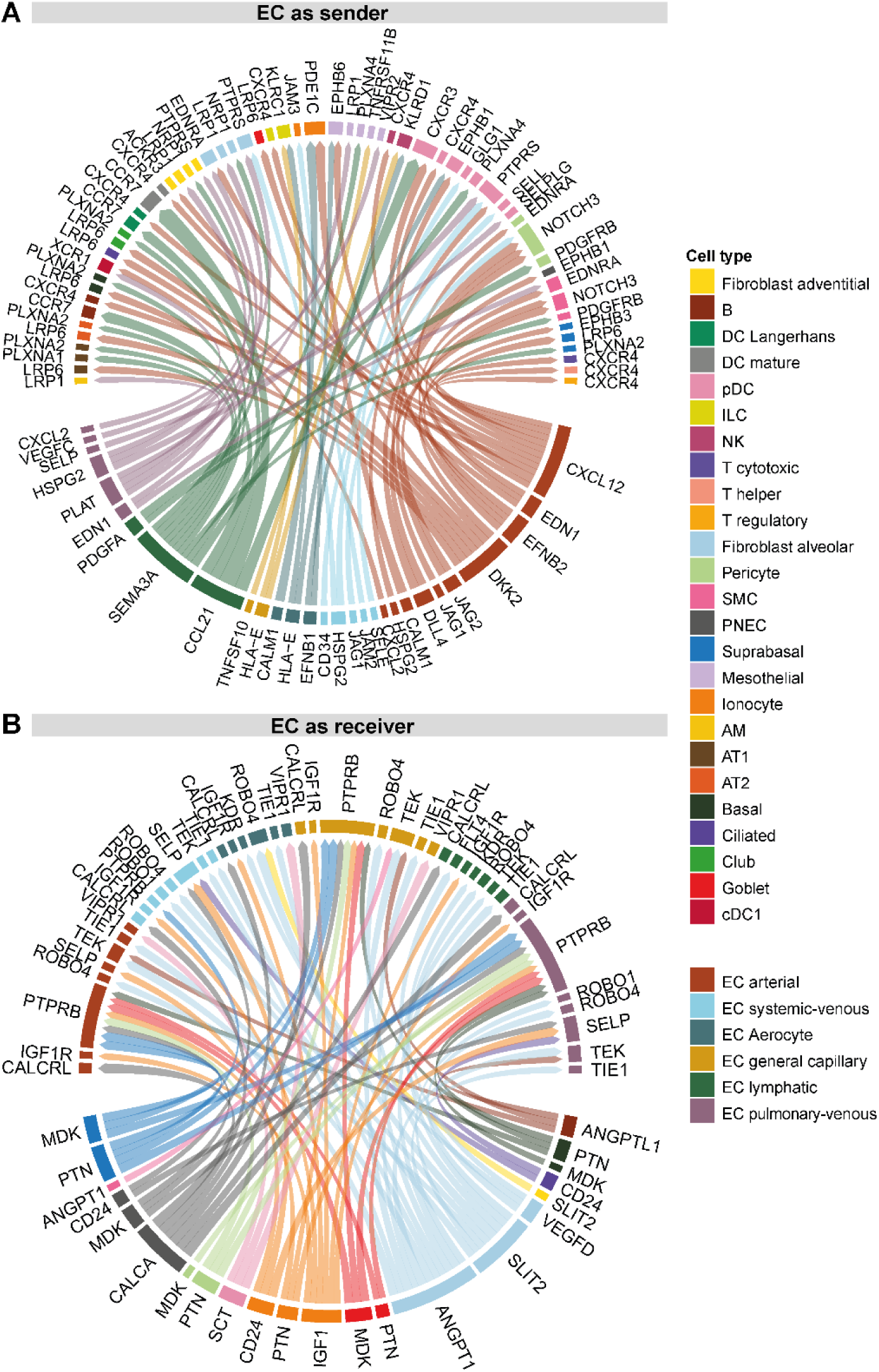
EC focused intercellular communication. Visualization of a small subset of the connectomic analysis. Circos plot of top 75 edges by edge weight with EC subpopulations as **(A)** “senders” and **(B)** as “receivers”. Edge thickness is proportional to edge weights. Edge color labels the source cell type. In both Circos plots ligands occupy the lower semicircle and corresponding receptors the upper semicircle. The full results of the connectomic analysis can be found in Table S6.

Lymphatic EC secrete CCL21, which is sensed by pDCs through CXCR3 and by mature DCs, Langerhans DCs and B-cells through CCR7 (Figure 4A), representing a classical mechanism for guiding matured DCs to secondary lymphoid tissues^41^. Lymphatic EC, as well as alveolar cell type 1 (AT1) and pericytes, express PDGFA, with the corresponding receptor PDGFRB expressed in pericytes and SMCs. Finally, lymphatic ECs were found to express the spatial guidance molecule SEMA3A that is critically important for lymphatic vessel maturation^14^. Its corresponding receptors are expressed on alveolar and non-parenchymal fibroblasts and vascular ECs (NPR1), AT1, AT2, Club, basal, suprabasal, and mesothelial cells (PLXNA1/PLXNA2), and pDCs (PLXNA4) (Figure 4A).

We next identified selected cell-cell signaling patterns with ECs being on the receiving end (Figure 4B). Alveolar fibroblasts strongly express the morphogen SLIT2 and ANGPT1 (to a lesser extent also in pericytes and SMCs). Both, SLIT2 and ANGPT1, can be sensed by all five EC subpopulations through the receptors ROBO4 and TEK/TIE1, respectively. Furthermore, alveolar fibroblasts are in a position to signal to aerocytes and lymphatic ECs through the VEGFD-KDR and VEGFD-FLT4 axes, respectively (Figure 4B).

It is worth noting that not all ligands sensed by EC receptors are produced by resident lung cells. For example, the receptor VIPR1 is expressed by lung ECs yet its main ligand VIP (Vasoactive Intestinal Peptide) was not expressed by any cells profiled in the lungs. This suggests that the VIPR1 ligands targeting lung ECs are mainly derived from extrapulmonary organs. Therefore, alternate sources of ligands beyond those profiled in the present connectomic analysis may contribute to lung EC signaling and function.

### Novel EC populations found in humans are also present in mice

In mammals, the lung is the only organ in which gas exchange between blood and air takes place. Here, we aimed to explore whether cellular and transcriptomic principles of lung vascular biology are preserved and chose the most used mammalian model organism as comparator: the mouse. We applied our human workflow to mouse lung scRNAseq data to generate an integrated dataset of mouse lung cells from 18 control wild-type C57BL/6 mice from six cohorts^12, 15, 37, 42-44^ (Supp. Figures S3, Supp. Table S7). All murine scRNAseq data had been generated from whole lung dissociations using the 10x 3prime scRNAseq kits based on V2 chemistry. Only 2 out of 18 mice were female and the median age was 12.5 weeks (IQR 10.75 – 34.0 weeks; for further details by cohort, see supplemental table S2).

We identified 21,831 ECs within the integrated mouse lung dataset containing 57,974 single cell transcriptomes (Figure 5A, Supp. Figures S3). Subsetting the ECs from the mouse lung dataset for more specific subtype identification revealed the same EC subpopulations found in humans (Figure 5A). Regarding cell type marker gene expression in the full mouse dataset and the EC subset, please refer to Supp. Figure S3&S4 and Supp. Table S8&S9 and to our data dissemination portal www.LungEndothelialCellAtlas.com, regarding technical summaries of sample preprocessing to Supp. Table S10.

**Figure 5.**
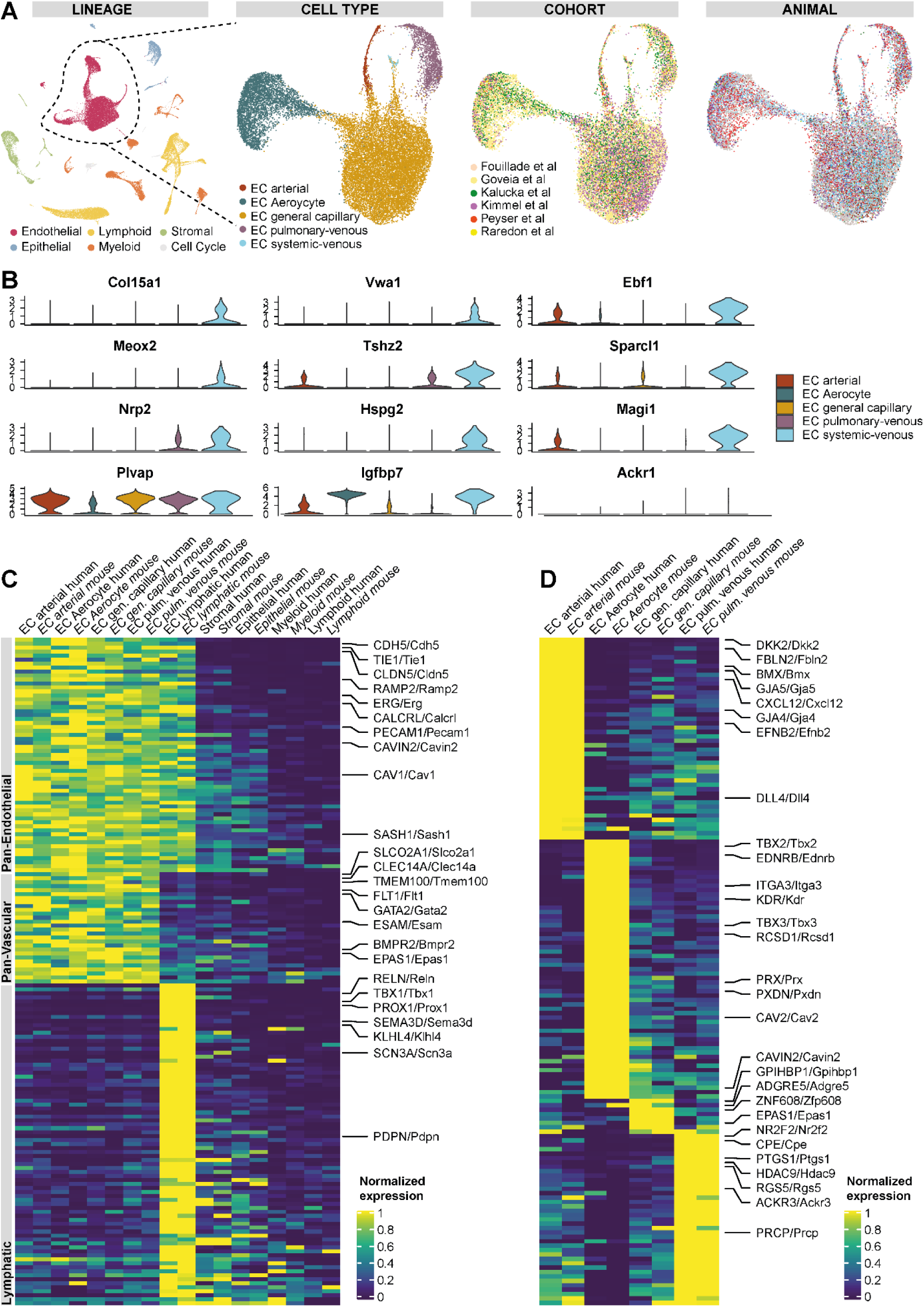
Identification of conserved EC populations and conserved marker genes in human and mice. **(A)** UMAPs of (i) all 57,974 mouse cells from 18 control mice lungs colored by lineage membership, and of the subset of 21,343 endothelial cells labeled by (ii) cell type, (iii) cohort and (iv) animal. In the UMAP colored by subjects, each color represents a distinct mouse. **(B)** Violin plots of marker genes significantly expressed in systemic-venous ECs compared to all other ECs (first till third row) and of homologues to human marker genes of systemic-venous ECs lacking specificity for murine systemic-venous ECs (fourth row) **(C)** Heat map of conserved pan-endothelial, pan-vascular and lymphatic EC marker genes. Each column represents the average expression value per cell type for ECs and per lineage for non-ECs. All gene expression values are unity normalized per species from 0 to 1 across rows. **(D)** Heat map of conserved marker genes of four pulmonary EC populations. Each column represents the average expression value per cell type. All gene expression values are unity normalized per species from 0 to 1 across rows. Human genes are indicated by capital letters and mouse genes are indicated by small letters after the first letter, separated by a slash. Labels of mouse, but not human, cell types / lineages are given in italics.

We identified the previously undescribed systemic-venous EC type in mice as well, based on the marker genes *Col15a1* and *Vwa1* (Figure 5B, Supp. Fig. 4A, Supp. Table S9). Systemic-venous ECs are very rare in mice and account for only 0.5% of all ECs (n=117 cells, Supp. Fig. 4B), probably because – unlike in humans - both the visceral pleura and intraparenchymal airways are supplied by the pulmonary rather than the systemic circulation^45, 46^. As in humans, murine systemic-venous ECs express transcription factors *Ebf1, Ebf3, Meox2* and *Tshz2*, phosphodiesterase *Pde7b* and, overlapping with venous ECs, *Selp* and *Ackr1*. However, homologues to some human marker genes of systemic-venous EC show no specificity for murine systemic-venous ECs (*Plvap, Spry1, Pkp4*), a different specificity profile (*Igfbp7*), a very low expression (*Tacr1, Robo1*) (Figure 5B), or a homologue simply doesn’t exist in mice (e.g. ZNF385D).

Murine capillary ECs are characterized by the expression of canonical markers *Car4*^12, 23^, *Prx*^24^, *Sgk1*^12^, and split likewise in two subpopulations: aerocytes (n= 4,853 cells) and general capillary ECs (n= 14,119 cells; Supp. Figure 4A, Supp. Table S8&9). Aerocytes were identified by the expression of *Ednrb* and *Tbx2* and general capillary ECs by *Gpihbp1* and *Edn1* (Supp. Table S9). In the median per mouse, capillary ECs were profiled in a ratio of aerocytes to general capillary ECs of 0.24:1 (IQR 0.15 - 0.39, Supp. Figure 2B) indicating that aerocytes are less common in mice lung capillaries when compared to those of humans (ratio in human of 0.48:1, IQR 0.26 – 1.16). A breakdown of marker genes overlapping with humans for both populations is given in the following section. In contrast to what’s observed in humans, murine aerocytes are additionally characterized by top marker genes like *Igfbp7, Emp2, Rgs6, Abcc3, Ackr2, Fibin, Chst1* and general capillary ECs by genes such as *Glp1r, Adgrl3, Plcb1* and *Hmcn1* (Supp. Table S9). Interesting patterns can be observed, which distinguish both murine capillary cell types, but which are only vaguely recognizable in humans: *Apln* is specifically expressed in aerocytes, while its receptor *Aplnr* is expressed in general capillary ECs. In contrast, *Vegfa* is expressed in general capillary ECs and its receptor *Kdr* on aerocytes (Supp. Table S9). In contrast to humans, major constituents of Weibel-Palade bodies - *Vwf* and *Selp* – are neither expressed in aerocytes nor in general capillary ECs.

### Endothelial marker genes are widely conserved in mice

We performed an unsupervised integration of data from both human and mice cells into a single graph embedding and subsequent UMAP plot (Supp. Figure S5). We observed the corresponding overlap of all five homologous EC subpopulations and, as a result, performed an in-depth comparison of corresponding populations between the two species. An analysis of the homologous marker genes across human and mouse EC subpopulations revealed a set of conserved marker genes between analogous subpopulations. In the following paragraph, human genes (all capital letters) and mouse genes (italicized) are separated by a slash.

Unsurprisingly, conserved marker genes expressed in all EC populations included canonical markers PECAM1/*Pecam1*, CDH5/*Cdh5* and CLDN5/*Cldn5* (Figure 5C, Supp. Table S11). Furthermore, all ECs from both species express the transcription factor ERG/*Erg* and SOX18/*Sox18*, angiopoietin receptor TIE1/*Tie1*, and the adrenomedullin receptor subunits CALCRL/*Calcrl* and RAMP2/*Ramp2*. Conserved vascular marker genes - genes not expressed in lymphatic ECs - are comprised of endothelial receptors (FLT1/*Flt1*, ADGRL4/*Adgrl4*, BMPR2/*Bmpr2*, ACVRL1/*Acvrl1)*, cell adhesion molecules (ESAM/*Esam*, CLEC14A/*Clec14a*), transcription factors (GATA2/*Gata2*, SOX7/*Sox7*) and a prostaglandin transporter (SLCO2A1/*Slco2a1*). Lymphatic ECs represent a distinct population in both species and are characterized by canonical PROX1/*Prox1*, semaphorin SEMD3D/*Sema3d*, the transcription factor TBX1/*Tbx1*, Podoplanin (PDPN/*Pdpn*) and matrix protease Reelin (RELN/*Reln*) (Figure 5C, Supp. Table S11).

Finally, we examined the conserved marker genes that are used to distinguish between lung EC subpopulations (Figure 5D, Supp. Table S11). Arterial ECs share a unique set of genes between both species that includes connexins GJA4/*Gja4* and GJA5/*Gja5*, secreted cytokines CXCL12/*Cxcl12* and EFNB2/*Efnb2*, Wnt modulator DKK2/*Dkk2* and Notch ligand DLL4/*Dll4*. Aerocytes are characterized by the conserved expression of receptors EDNRB/*Ednrb* and KDR/*Kdr*, transcription factors TBX2/*Tbx2* and TBX3/*Tbx3* and peroxidase PXDN/*Pxdn* (Figure 5D, Supp. Table S11). Pulmonary-venous EC show conserved expression of transcription factor NR2F2/*Nr2f2*, receptors IL1R1/*Il1r1 and* ACKR3/*Ackr3, c*arboxypeptidases CPE/*Cpe* and PRCP/*Prcp*, and the marker genes HDAC9/*Hdac9*, PTGS1/*Ptgs1*, SELP/*Selp* and RGS5/*Rgs5*. The least overlapping marker genes were observed in general capillary ECs, which included GPIHBP1/*Gpihbp1*, EPAS1/*Epas1*, ZNF608/*Zfp608*, ADGRE5/*Adgre5* and NTRK2/*Ntrk2* in both species (Figure 5D, Supp. Table S11).

## Discussion

In this study, we present a reference atlas of endothelial cells of the human lung on a cellular transcriptomic level derived from 15,142 endothelial cells from 73 subjects. We identified prototypic gene expression patterns specific to distinct hierarchies of the endothelial lineage including pan-endothelial, pan-vascular and subpopulation-specific marker gene sets. Importantly, the expression of all reported marker sets was detected throughout all cohorts and most subjects, essentially eliminating the possibility that subject-specific gene expression patterns could be misattributed as cell type specific patterns. We validated major marker genes on the protein level by immunohistochemistry, or at the mRNA level by in-situ hybridization, respectively. We uncoupled the identities of two previously indistinguishable EC populations: pulmonary-venous ECs localized to the lung parenchyma, and systemic-venous ECs localized to the airways and the visceral pleura. We confirmed and characterized the new subclassification of pulmonary capillary ECs into aerocytes and general capillary ECs. We investigated the role of EC subpopulations within the complex cellular signaling networks that occur between different lung cell types, emphasizing the essential supporting role of alveolar fibroblasts with respect to all lung ECs though the ANGPT1-TEK/TIE1 and SLIT2-ROBO4 axes, and the crucial arterial ECs to mural pericytes/SMCs communication by the supportive DLL4/JAG1/JAG2-NOTCH3 axis^38^ and the vasoconstrictive EDN1-EDNRA axis^39^.

The most significant contribution of this study represents the unbiased and global-transcriptomic characterization of EC of the human lung on three hierarchic levels: we identified 147 genes specifically expressed in all EC populations, 142 genes specifically expressed in vascular ECs, and describe in-depth the transcriptional particularities of the six endothelial cell types. Pan-endothelial, pan-vascular or EC subpopulation specific gene expression was often associated with a specific function of endothelial cells in general or of endothelial subpopulations in particular. This is exemplified in the expression of genes associated with immune cell homing in lymphatic ECs, with gas exchange in capillary ECs, with diapedesis of leukocytes and unspecific transcytosis of chemokines in venous ECs or with ECM components of the elastic vessel wall and regulation of vascular tone in arterial ECs. However, the function of many other genes specifically expressed in ECs is not known, and we hope that their future study will greatly enhance our understanding of endothelial cell biology.

Our integrated analysis confirmed and expanded the description of two novel EC cell subpopulations. We had previously noticed a distinct population of COL15A1 ^pos^ lung ECs that seemed to appear in the lung parenchyma only in IPF lung^10^. Here, basing our analysis on many more healthy lungs, we identify this population as a venous EC population that, as revealed through in-depth immuno-histological studies, is usually located in the peribronchial space and the visceral pleural. This discovery is an example of the power of unbiased scRNAseq profiling, as it is hard to think this distinct population could be identified using traditional methods. Whether the specific gene expression profile of systemic-venous ECs reflects adaptation to systemic perfusion of blood vessel of bronchi and the visceral pleura or to the extracellular matrix itself in which they are located will need to be investigated in future studies, but the fact that this EC subtype changes its anatomical distribution in pulmonary fibrosis may suggests involvement in repair and relevance to human lung disease.

Similarly, studies in multiple species on pulmonary scRNAseq datasets identified two distinguishable capillary EC populations^10, 12, 15, 34, 37, 47, 48^. This consistent, but unexpected observation, for which there is no corresponding report from the pre-scRNAseq era, provides an explanation for the mosaic or patchy immuno-histological staining patterns of VWF, THBD, EMCN, EDN1 and EDNRB^15, 28-33^ in lung capillaries that had, until recently, remained enigmatic. One of these capillary EC has been recently defined as aerocytes^34^: the key endothelial cells involved in gas exchange. Intriguingly, aerocytes – unlike all other vascular EC types – lack expression of the major constituents of endothelial-specific Weibel-Palade bodies including VWF, SELP, EDN. Weibel-Palade bodies are known to be missing in the thinnest lung capillaries, which are thought to be the major location were gas exchange happens^49, 50^. And indeed, Vila Ellis et al. and Gillich et al. showed by lineage-tracing experiments in mice that aerocytes exhibit a much larger surface area and form - together with the AT1 cells and their interposed basement membrane - the blood-air-barrier^34, 47^. In addition to confirming their presence in large numbers of individuals and in mice, our study adds in-depth transcriptional profiling of aerocytes, thus providing additional clues of their potential functions.

The comparison of human and mouse EC diversity and marker gene expression revealed considerable overlap. All human EC subpopulations could be identified in mice as well, including a very rare systemic-venous *Col15a1*^pos^ EC population. The rarity of these systemic-venous ECs is probably due to the fact that the systemic bronchial circulation in mice is very limited and reaches only the main bronchi^46^. Several dozens of pan-endothelial and pan-vascular marker genes and of marker genes specific to the EC subpopulations are conserved in mice and human. Marker genes of general capillary ECs are an exception as they show only a limited overlap, potentially due to fewer marker genes per species in general and a lack of homologues to some human marker genes (e.g. FCN3) in mice.

Our study is not without limitations: First, in order to enable a comprehensive joint analysis of all datasets, a data integration approach had to be chosen (details see supplement). Each cohort entails a batch effect due to slightly different processing of samples and variation in the single cell RNA seq library preparation and sequencing. A major technical batch effect was alleviated by processing the raw sequencing data through the same computational pipeline and the same reference genome. Importantly, all differential testing was performed on the non-integrated data to ensure that it was not distorted by the integration algorithm. However, integration of several datasets has the advantage of leveling out the cell preparation/dissociation bias, i.e. the preferential liberation of specific cell types during tissue dissociation. Second, although we validated and localized the expression of major endothelial marker genes on the protein level by immunohistochemistry and on the mRNA level by in situ hybridization, a global spatial localization of endothelial gene expression is lacking in this report. We assume that recent advances in spatial transcriptomics will enable this in the future. Third, the connectomic analysis relies on a limited database of known ligand-receptor interactions and can only infer potential communication between cell types. A sensible future approach would be to study the molecular crosstalk in functional studies.

The generation of transcriptional profiles of cell types and their intercellular communication within the complex organ lung are tasks that are crucial to understanding the molecular function of the lung, and its dysfunction in disease. We hope that this integrated lung endothelial atlas will advance lung endothelial research, especially in diseases having vascular involvement. For this purpose, we have created the online tool www.LungEndothelialCellAtlas.com with which all transcriptomic data of endothelial cells of the lung can be explored in an easily accessible fashion.

## Acknowledgements

We thank all donors who participated in the several studies analyzed in this manuscript, all funders of these studies, and the authors who made these rich datasets publicly available. We thank M. Zhong at Yale Stem Cell Center for the sequencing service and A. Brooks and his team at Yale Pathology Tissue Services for tissue processing.

## Author’s contributions

JCS and NK conceptualized and supervised the study. JCS, TSA and SP procured and dissociated the lungs. JCS and TSA performed scRNAseq barcoding and library construction. Data were processed, curated, integrated and visualized by JCS, TSA and XY and analyzed by JCS, TSA, MSBR, XY, NK. The online tool “www.LungEndothelialCellAtlas.com” was developed by CC and NN. IHC staining was performed by JCS and KAR, RNA in situ hybridization staining by NO, both of which were evaluated by RH, NK, JCS and NO. The manuscript was drafted by JCS, TSA, CC, MSBR and NK and was reviewed and edited by all other authors. MSBR, LEN, YY, ACH, AJG, LTB, SAT, MCN, KBM, DP, NEB, JAK provided datasets and insights, LEN, SAT, MCN, NEB, JAK, IOR, NK acquired funding for the integrated datasets.

