## Supplement for "Integrated Single Cell Atlas of Endothelial Cells of the Human Lung"

#### Methods:

Sample preparation, barcoding and library preparation for single-cell sequencing

While most data analyzed herein is derived from previously published datasets, we also subjected four additional control lung samples to single cell RNA sequencing (scRNAseq). These new samples were procured and processed similarly to the control samples from our previous publication[1]. Healthy lungs were rejected donor organs that underwent lung transplantation at the Brigham and Women's Hospital, or donor organs provided by the National Disease Research Interchange (NDRI). The study protocol was approved by the Partners Healthcare Institutional Board Review (IRB Protocol # 2011P002419) and the Yale University Institutional Review Board (IRB Protocol ID: 2000022618). Briefly, lung specimens were enzymatically dissociated. After several washing steps and red cell lysis, cell suspensions were frozen and stored in liquid nitrogen till further processing. After thawing, cells were filtered and washed again, then processed using the 10x Chromium single cell RNA seq platform (Single Cell 3′ Reagent Kits v2, 10× Genomics, USA). Single-cell barcoding and library preparation was performed according to the manufacturer's protocol (Single Cell 3′ Reagent Kits v2, 10× Genomics, USA) with a targeted cell output of 10,000 cells per library. cDNA and final libraries were analyzed on an Agilent Bioanalyzer High Sensitivity DNA chip for qualitative control purposes. cDNA libraries were sequenced with a HiSeq4000 Illumina aiming for 150 million reads per library with a sequencing configuration of 26 base pairs on read1 and 98 base pairs on read2. Basecalls were converted to reads using the software Cell Ranger (v3.0.2).

Read processing

The dataset repositories GEO, dbGaP, and EMBL-EBI were searched for scRNAseq datasets of human lungs generated using the 10x Single Cell RNAseq technology (V2 chemistry). Control samples of the following human lung scRNAseq datasets used (for overview of individual libraries, see supplemental table S1): EGAD00001005064/5[2] ("WSI Groningen"), phs001750.v1.p1[3] ("Northwestern"), GSE135893[4] ("Vanderbilt TGen"), E-MTAB-6149/6653[5] ("Leuven VIB"), E-MTAB-6308[6] ("Leuven LKI") and GSE136831&GSE133747[1, 7] ("Yale Baylor"). FASTQ files containing raw sequencing reads were either provided by the authors or downloaded from the appropriate repositories.

Furthermore, dataset repositories GEO and EMBL-EBI were searched for scRNAseq datasets of mouse experiments generated using the 10x Single Cell RNAseq technology (V2 chemistry) and were available without missing relevant sequencing data (i.e. without missing read1 or read2 data). Control samples of the following mouse lung scRNAseq datasets were identified and FASTQs downloaded from repositories (for overview of individual libraries, see supplemental table S7): E-MTAB-7458[6], E-MTAB-8077[8], GSE133747[7], GSE129605[9], GSE132901[10] and GSE133992[11].

Multiple FASTQs from the same library and read1/read2 were concatenated to single files. Adaptor contamination (AAGCAGTGGTATCAACGCAGAGTACATGGG 10x 3prime samples; CCCATATAAGAAA for 10x 5prime samples) and 20bps or longer poly(A) or poly(T) sequences (for samples from 10x 3prime or 5prime kits respectively) were removed using cutadapt (v2.9). After trimming, read pairs were removed if any of the reads was trimmed below 25 bp.

Subsequent read processing was conducted using the scRNAseq implementation STARsolo of the software STAR[12] (v2.7.3a). Reads were aligned to the human reference genome GRCh38 release 31 (GRCh38.p12) or the mouse reference genome GRCm38 release M22 (GRCm38.p6), both downloaded from GENCODE[13]. Collapsed unique molecular identifiers (UMIs) with reads that span both exonic and intronic sequences were retained as combined gene expression assays, as well as exonic-only derived UMIs gene expression assays. Valid cell barcodes representative of quality cells were distinguished from barcodes of dying cells or background RNA based on the following three thresholds, as described before[1]: at least 7.5% of transcripts arising from unspliced reads indicative of nascent mRNA; more than 1000 transcripts profiled; less than 20% of their transcriptome was of mitochondrial origin. Cell free mRNA background contamination was removed using the software SoupX (v1.2.2)[14]. Technical summaries related to sequencing and data processing can be found in supplemental table S4 for human samples and S10 for mouse samples.

<u>Dataset integration, endothelial cell identification</u>

UMIs from each valid cell barcode, irrespective of whether they derived from spliced or unspliced mRNA, were retained for all downstream analyses and analyzed using the R package Seurat (version 3.1.4)[15]. UMI counts were normalized with a scale factor of 10,000 UMIs per cell and then natural log transformed using a pseudocount of 1. In order to identify and isolate endothelial cells from cells that belong to other lineages, cells from all datasets were subject to a recursive process of integration, graph-embedding and cluster analysis. For each iteration, integration of datasets and clustering was performed as recommended in Seurat (https://satijalab.org/seurat/v3.0/integration.html). Briefly, the top variable genes within each dataset were selected using the Seurat implementation *FindVariableGenes* using the "vst" parameter. Shared patterns of variance in these genes within each dataset where then used to integrate the datasets using Seurat's *FindIntegrationAnchors* and *IntegrateData*[15], then the resulting integrated expression matrix was scaled using Seurat's *ScaleData*. These scaled values were then used for principal component analysis (PCA), and the top components were used for 2-dimensional Uniform Manifold Approximation and Projection (UMAP) graph embedding and Leiden clustering. Next, each unsupervised cluster was assessed for sample and dataset representation and within-cluster gene expression consistency. Clusters without noteworthy gene expression differences between them were collapsed. Clusters comprised of heterogeneous cell populations were subsetted alongside phenotypically similar clusters, then subject to another iteration of data integration and cluster analysis. This process continued until all cells from all datasets could be assigned to a discrete cell type, whose distinguishing features are consistently represented across different subjects and datasets. After comprehensively categorizing all cells into distinct cell-type clusters, we then determined each cluster's respective lineage assignment based on its expression profile of classical lineage markers. Five clusters were found to co-express classical endothelial markers PECAM1 and CDH5; notably these clusters were the only clusters collectively missing expression of classical markers of epithelial (EPCAM, CDH1), mesenchymal (PDGFRA, PDGFRB) and immune cells (PTPRC). Mesothelial cells were grouped with mesenchymal cell types for reasons of simplicity. As lymphatic ECs were substantially different from vascular ECs, establishing marker genes that enable differentiation of subvarieties within vascular ECs deemed most important. Therefore, a final subset consisting of only vascular ECs was created for downstream analyses. This final vascular EC subset also included a sixth dataset ("Leuven LKI") containing sorted pulmonary EC cells. As two samples of the total of 75 subjects did not contain any ECs, the vascular ECs is derived from 73 subjects only. Sample "NEC50" of the cohort "Leuven LKI" was found to contain relevant cluster of cancer cells (n=789) and therefore not included in this study.

Multiplets and other cell barcodes of low quality were identified using a multilayer approach: First, the software DoubletFinder[16] was used to predict multiplet clusters in an automated and unbiased fashion using an estimated multiplet rate of 0.8% per 1000 cells, according to 10x Genomics Single Cell 3′ Reagent Kits v2 protocol. DoubletFinder was applied per sample due to very different cell numbers per sample and, by that, different expected multiplet rates. However, DoubletFinder was not able to identify several multiplet clusters, especially of cell types with low frequencies. Therefore, multiplet clusters were additionally identified manually as having a transcriptomic signature that resembled the combination of two or more different cell type signatures that already existed in the data set. This approach was applied to the full dataset as well as all lineage subsets. Barcodes identified as multiplets or being of low quality were not included in any downstream analyses.

<u>Identification of cell type specific marker genes</u>

Cell type specific marker genes were identified using the Wilcoxon rank sum test by comparing all cells within a specific cluster to all cells outside said cluster. All p-values were adjusted for multiple comparisons using the conservative Bonferroni correction. Pan-endothelial marker genes were defined as genes significantly expressed in all bona fide pulmonary EC populations (arterial, pulmonary-venous, aerocytes, general capillary ECs, incl. lymphatic ECs) and a logFC of greater than 0.25, when compared to all other cell types. Pan-vascular marker genes were defined as genes significantly expressed in all the bona fide pulmonary vascular EC populations (arterial, pulmonary-venous, aerocytes, general capillary ECs) and a logFC of greater than 0.25, when compared to all other cell types, but not in lymphatic ECs.

As described before[1], we used an additional approach of a binary classifier system to assess the utility of detecting a given gene for classifying a cell. For each specific cell type, we selected all genes whose expression was log fold change ≥ 0.25 greater in all other cells in the data. Of those genes, we calculated the diagnostics odds ratio (DOR), where we binarize the expression values by treating any detection of a gene (normalized expression value > 0) as a positive value and zero expression detection as negative. To avoid undefined values, we included a pseudocount of 0.5 as follows:

$$DOR = ((TruePositives+0.5)/(FalsePositives+0.5))/((FalseNegatives+0.5)/(TrueNegatives+0.5))$$

where TruePositives represents the number of cells within a cluster detected expressing the gene (value > 0), FalsePositives represents the number of cells outside of the cluster detected expressing the gene, FalseNegatives represents the number of cells within the cluster with no detected expression, and TrueNegatives represents the number of cells outside of the cluster with no detected expression of the gene. Log-transformed DOR values are given in all supplemental tables on marker genes.

Connectome Analysis:

To study cell-cell signaling, the data was mapped to the NicheNet ligand-receptor interaction database[17] using the R software Connectome (v0.2.2) (https://msraredon.github.io/Connectome/)[7]. In brief, each cell type was treated as a single node for network creation. Average expression values, for all data slots, were calculated for all ligand and receptor genes on a per-cell-type basis, and an unfiltered edgelist ("connectome") was created linking all producers of a ligand to all producers of a receptor, with associated quantitative edge attributes. This mapping leveraged only well-annotated and literature supported ligand-receptor interactions (i.e., "kegg_cytokines", "kegg_cams", "kegg_neuroactive", "kegg_ecm", "pharmacology", "ramilowski_known"). Selected ligand-receptor interactions were visualized as Circos plots using the R package *circlize*[18] after filtering the connectome based on the following criteria: Ligands being significantly expressed in EC subpopulations and receptors significantly expressed in non-EC populations or vice versa (p<1e-5 based on a system-wide Wilcoxon Rank Sum test); ligands and receptors being expressed in at least 20% of the cells of their respective cell types with ligand and receptor mean scaled expression values > 0; omitting all integrin receptors due to their promiscuity; and finally, selecting the top 75 interactions for both Circos Plots in Figure 4, when ranked based on the scaled weight, defined as the mean of the scaled expression values of the ligand and receptor.

Analysis of mouse datasets

Integration, clustering, cell type annotation and multiplet identification of control mouse datasets was performed as described above for the human samples. When using the Seurat implementation *FindIntegrationAnchors* to integrate the mouse dataset, we reduced the default parameter for neighbors for filtering from 200 to 150, in order to accommodate a dataset with less than 200 cells, with otherwise default settings. Cell were assigned to lineages based on canonical marker genes as follows: lymphoid (Ptprc+ and Cd79a+ or Cd2+), myeloid (Ptprc+ and Lyz2+), endothelial (Cldn5+), epithelial (Epcam+) and stromal (Pdgfra+ or Pdgfrb+) lineages. Cell type annotations were performed to a similar granularity compared to humans, e.g. cDC2a and cDC2b were kept as cDC2. Barcodes identified as either multiplet or low quality were not included in any downstream analyses. Identification of cell type marker genes were performed as described above for the human samples.

Comparison of mouse and human ECs

For comparison of EC marker gene expression between human and mouse, mice gene names were translated using the R package biomaRt[19] if there was a one-to-one homologue available. As a proof-of-principle, we performed an integration of mouse and human control ECs. To this end, both barcode-gene-matrices were subsetted, keeping only genes for which a one-to-one homologue in the respective other species was available. All datasets were integrated following the same integration workflow described for the separate human and mouse analyses above. The only parameter change made was a change from 200 to 150 neighbors for anchor filtering with the Seurat package's *FindIntegrationAnchors* implementation, for the same reasons described in the mouse analysis methods, above.

To identify conserved pan-endothelial, pan-vascular or specific marker genes of the bona fide pulmonary vascular ECs (aerocytes, general capillary, arterial, pulmonary-venous) and lymphatic ECs, datasets were randomly downsampled to 500 cells per cell type in both species, to ensure balanced proportions. Identification of cell type specific marker genes of this balanced dataset was performed using the Wilcoxon Rank sum test with adjustment for multiple comparisons as described above. Conserved marker genes are defined as being uniquely found differentially expressed in the same cell type of both species. Conserved lymphatic marker genes were defined as genes being significantly expressed in lymphatic ECs compared to all other lung cells in both species. Conserved pan-endothelial marker genes were defined as genes being significantly expressed in all four bona fide pulmonary vascular and lymphatic ECs compared to all other lung cells in both species. Conserved pan-vascular marker genes were defined as genes being significantly expressed in all four bona fide pulmonary vascular ECs when compared to all other lung cells in both species, but not significantly expressed in lymphatic ECs compared to all other lung cells in both species. For results of this comparison between mouse and human ECs, please refer to table S11.

Immunohistochemistry

Immunohistochemistry was performed as previously described[1]. FFPE (Formalin fixed paraffin embedded) blocks were cut at 5 µm, rehydrated (xylene/ethanol deparaffinization), then boiled at 95°C for 20 min in 1× Tris-based Antigen Unmasking Solution (Vector Laboratories, USA) for heat-induced antigen retrieval. Histology slides were incubated for 10 min in BLOXALL Blocking Solution (Vector Laboratories, USA) to block endogenous peroxidase and alkaline phosphatase activity. Unspecific antibody binding was blocked using 2.5% Normal Horse Serum Blocking Solution (Vector Laboratories, USA) for 20 min. Slides were incubated with the primary antibody (rabbit antibodies: ACKR1 (polyclonal, # PA5-82549, Thermofisher), CA4 (clone #039, # 10472-R039-50, Sino Biological), PRX (polyclonal, # NBP1-89598-25ul, Novus Bio), COL15A1 (polyclonal, # PA553667, Thermofisher), VWA1 (polyclonal, # 14322-1-AP, Proteintech); mouse antibodies: PECAM1 (clone JC/70A, #MA513188, Thermofisher), CLDN5 (clone A-12, # sc-374221, Santa Cruz), GJA5 (clone B-3, # sc-365107, Santa Cruz), PDPN (clone D2-40, # 916601, Biolegend); goat antibody: LYVE1 (polyclonal , # AF2089, Novus Bio), diluted in 2.5% Normal Horse Serum Blocking Solution for 30 min at room temperature. Histology slides were incubated for 30 min with secondary antibodies (anti-mouse/-rabbit/-goat ImmPRESS reagent, Vector Laboratories, USA), conjugated with horseradish peroxidase. Specimen were incubated for 10 min in DAB working solution (Vector Laboratories, USA). Slides were counterstained in Hematoxilin Solution Gill no. 1 (Sigma-Aldrich, USA) for 3 min, and then washed with tap water. Slides were dehydrated in ethanol/xylene and mounted with VectaMount permanent mounting solution (Vector Laboratories, USA). Stained slides were digitalized on an Aperio Scanner (Leica) and then analyzed using the softwares QuPath and ImageJ.

RNA in situ hybridization

RNAscope technology (Advanced Cell Diagnostics (ACD), Newark, CA) was used for RNA in situ hybridization (RNA-ISH). FFPE blocks were cut at 5 µm, mounted on slides, baked for 1 h at 60°C, then deparaffinized in xylene and 100% ethanol. Hydrogen peroxide (ACD 322381) was applied for 10 min at room temperature, followed by a mild boil at 98-102°C for 15 min in 1x target retrieval reagent buffer (ACD 322001). Sections were treated with Protease Plus (ACD 322381) at 40°C for 30 min in HybEZ Oven (ACD). Hybridization with target probes (FCN3 - ACD 818741, SOSTDC1 - ACD 469921), preamplifier, amplifier, label and wash buffer (ACD 320058) were performed following the ACD manual. Parallel sections were incubated with ACD positive (HsPPIB; ACD 313901, MnPpib; ACD 313911) and negative (DapB; ACD 310043) control probes.

### Supplemental Figures

#### Supp. Fig. S1

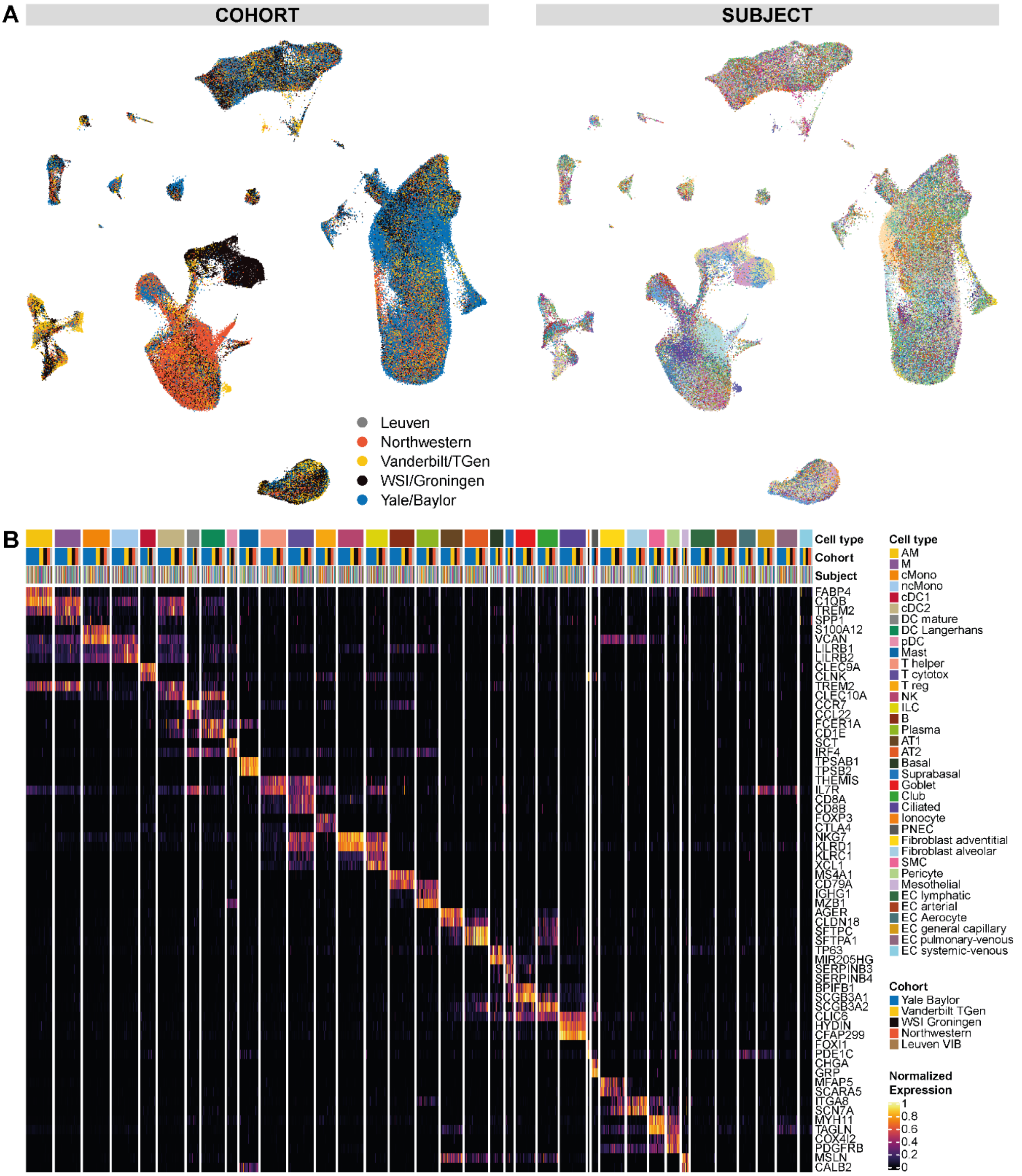


Supp. Fig. S1: **(A)** Additional annotations of the UMAP in Fig. 1B in the main manuscript. UMAP of the full dataset colored by (i) cohort and (ii) subjects. In the UMAP colored by subjects, each color represents a distinct subject. **(B)** Enlarged heat map of Fig. 3A zooming in on marker gene expression of all non-EC populations. Each column represents the average expression value for one subject, grouped by cell type and cohort. All gene expression values are unity normalized from 0 to 1 across rows.

**Supp. Fig. S2**

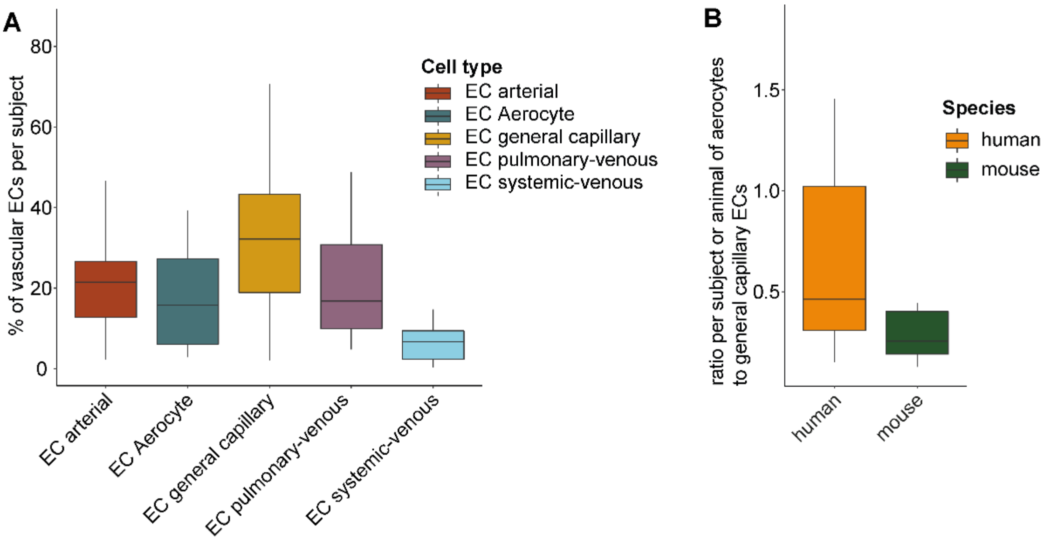


Supp. Fig. S2: **(A)** Boxplots representing the percentage of each vascular EC types among all vascular ECs per subject for all subjects in which all five vascular ECs had been profiled (n=29 subjects). The length of each whisker represents 1.5×IQR. **(B)** Boxplots representing the ratio of aerocyte to general capillary EC counts per human subject in which both cell types had been profiled (n=46 subjects) in orange and per mouse in green. The length of each whisker represents 1.5×IQR.

#### Supp. Fig. S3

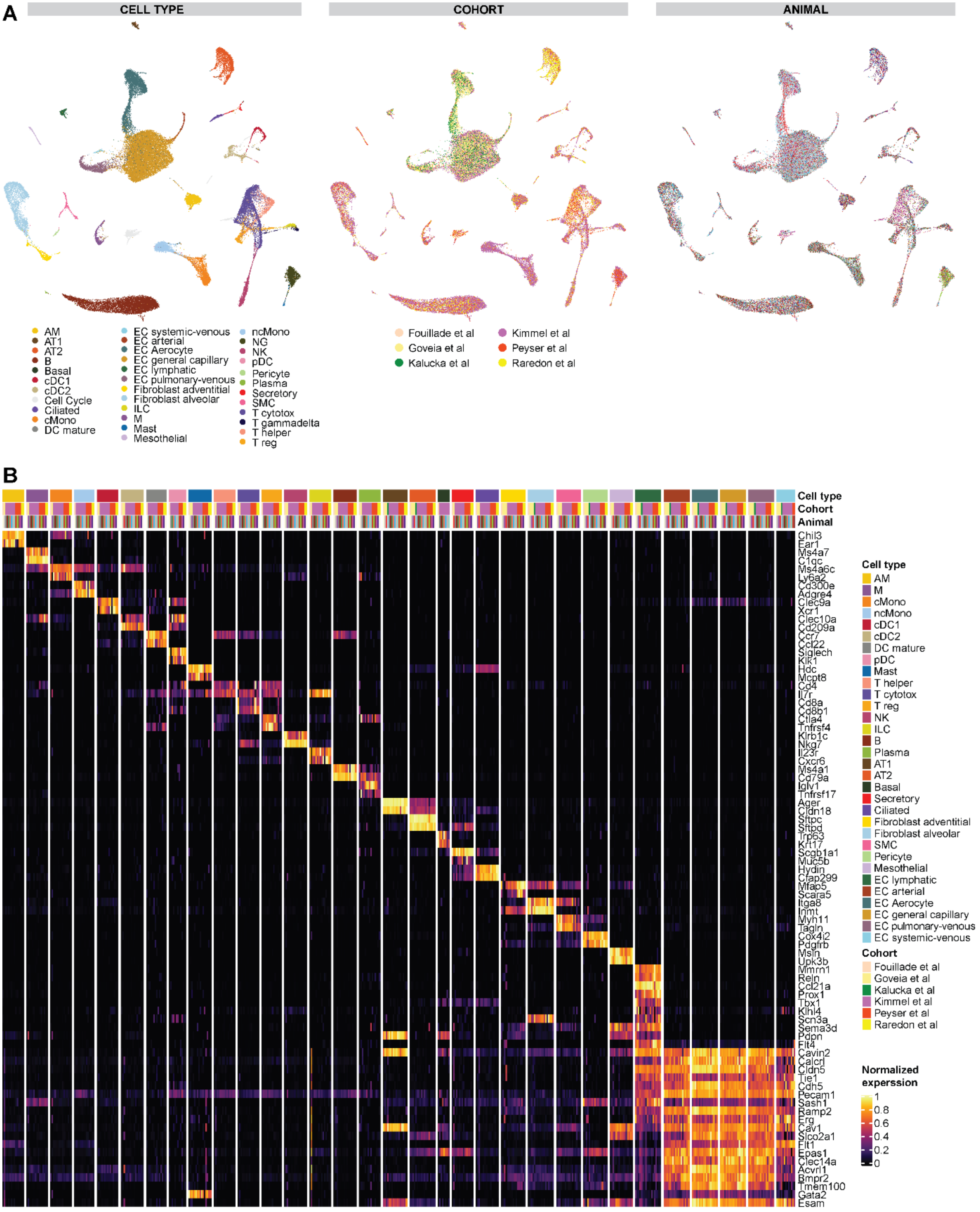


Supp. Fig. S3: **(A)** Additional annotations of the UMAP in Fig. 5A in the main manuscript. UMAPs of the full mouse dataset colored by (i) cell type, (ii) cohort and (iii) animals. In the UMAP colored by animals, each color represents a distinct library. **(B)** Heat map of murine marker genes for all non-EC cell types and of lymphatic cells, as well as pan-endothelial (specifically expressed in all EC sub-populations) and pan-vascular (specifically expressed in all EC populations but lymphatic ECs) marker genes. Each column represents the average expression value for one animal, grouped by cell type and cohort. All gene expression values are unity normalized from 0 to 1 across rows.

**Supp. Fig. S4**

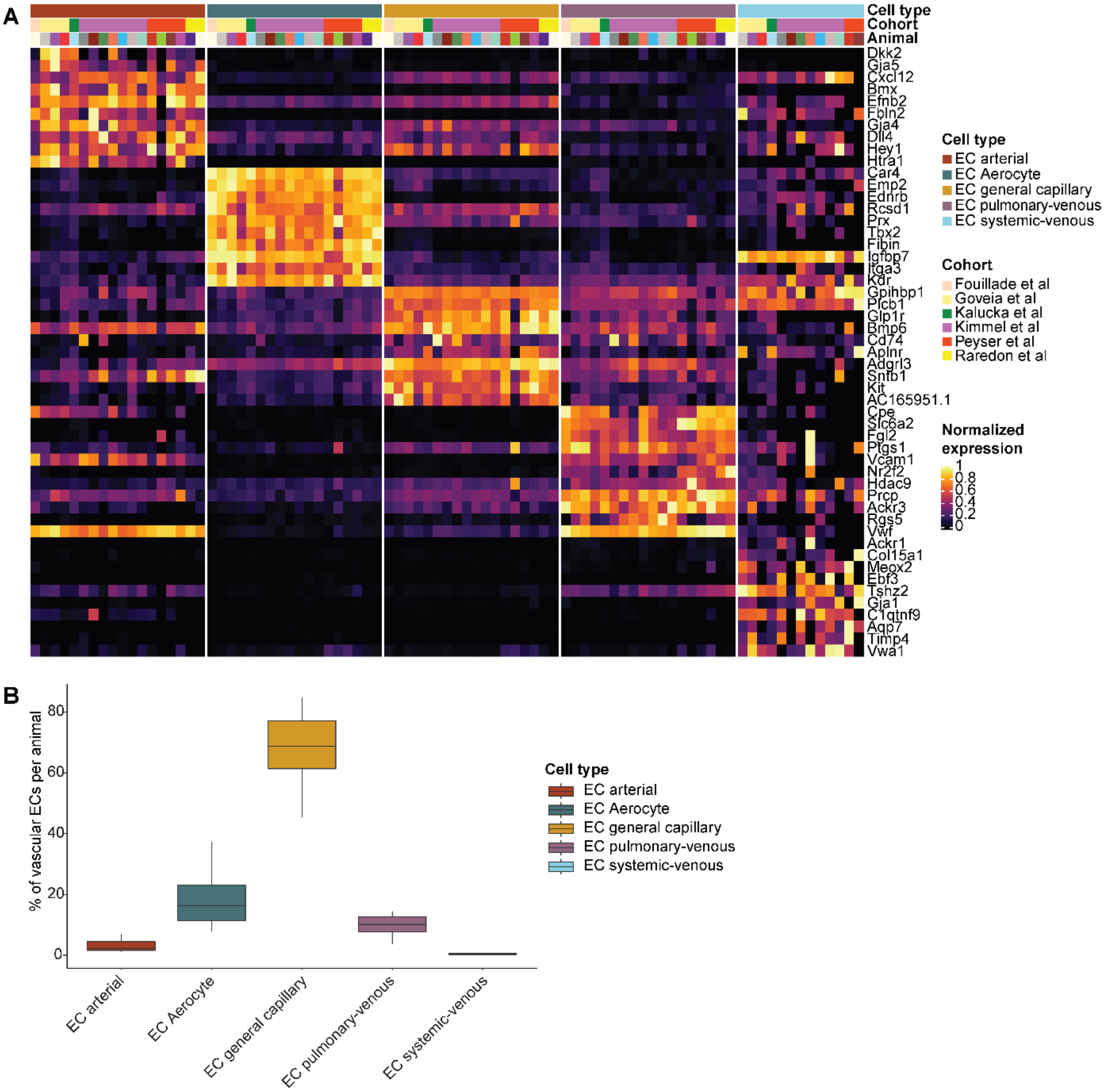


Supp. Fig. S4: **(A)** Heat map of marker genes all five EC populations of the mouse dataset. Each column represents the average expression value for one animal, grouped by cell type and cohort. All gene expression values are unity normalized from 0 to 1 across rows. **(B)** Boxplots representing the percentage of each murine vascular EC types among all murine vascular ECs per animal. The length of each whisker represents 1.5×IQR.

**Supp. Fig. S5**

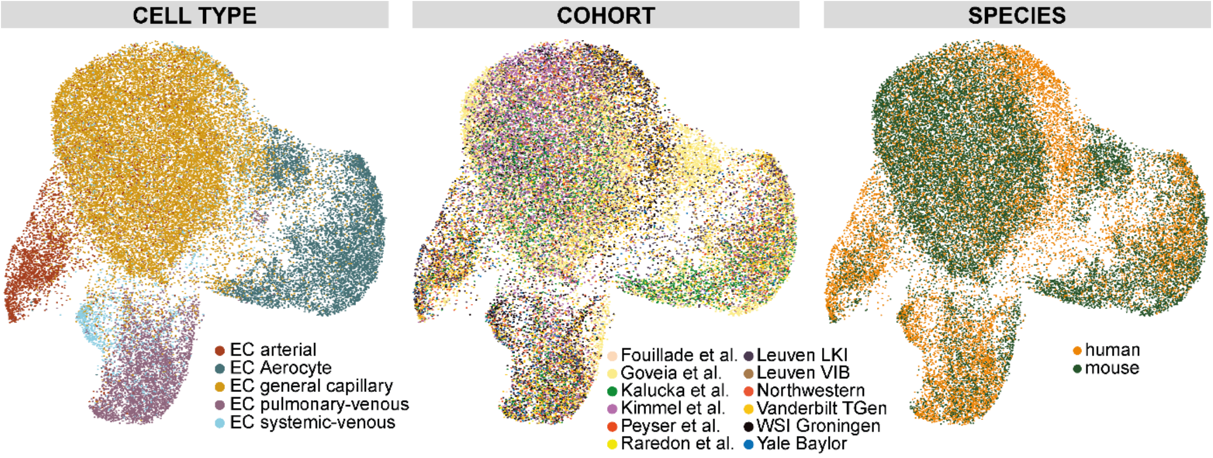


Supp. Fig. S5: UMAPs of integrated human and mice EC datasets. Each dot represents a single cell, and cells are labelled – from left to right - by (i) cell type, (ii) cohort and (iii) species.

#### Supplemental Tables

**Supp. Table S1: Table of libraries of all human data sets**
Table on basic characteristics of all human libraries included in this study.

**Supp. Table S2: Summary table of basic characteristics by cohort for human and mice**
Summary table on basic characteristics by cohort for human subjects and mice. If age was given as range in the original publication, the range was averaged for this purpose here. Age is given with interquartile range in square brackets. If the sums of certain variables do not add up to the total number of subjects per cohort, the corresponding information was not available for respective subjects. NA: not available; n. a.: not applicable.

**Supp. Table S3: Human cell type marker table and table of pan-endothelial and pan-vascular marker genes**
Results of Wilcoxon rank-sum test testing each cell type against all others. Each gene is annotated whether it was identified as pan-endothelial or pan-vascular marker gene. "pct.1" represents the fraction of cells within a specific cell type expressing a specific gene; "pct.2" represents the fraction of cells outside a specific cell type expressing a specific gene.

**Supp. Table S4: Technical summaries of library read processing of the human samples**
Technical summaries of computational read processing pipeline for each library.

**Supp. Table S5: Human vascular endothelial subset marker table**
Results of Wilcoxon rank-sum test of each vascular endothelial cell type against the other vascular endothelial varieties. "pct.1" represents the fraction of cells within a specific cell type expressing a specific gene; "pct.2" represents the fraction of cells outside a specific cell type expressing a specific gene.

**Supp. Table S6: Connectomic edgelist**
Connectomic edgelist generated on the basis of the full human dataset, filtered such that 20% cells within a given cell type express a specific ligand or receptor.

**Supp. Table S7: Summary table of libraries of all murine data sets**
Summary table on basic characteristics of animals and basic information of data sets for all murine libraries included in this study.

**Supp. Table S8: Mouse cell type marker table and table of pan-endothelial and pan-vascular marker genes**
Results of Wilcoxon rank-sum test of the mouse full dataset testing each EC type against all others. Each gene is annotated whether it was identified as pan-endothelial or pan-vascular marker gene. "pct.1" represents the fraction of cells within a specific cell type expressing a specific gene; "pct.2" represents the fraction of cells outside a specific cell type expressing a specific gene.

**Supp. Table S9: Mouse vascular endothelial subset marker table**
Results of Wilcoxon rank-sum test of each mouse vascular endothelial cell type against the other mouse vascular endothelial varieties. "pct.1" represents the fraction of cells within a specific cell type expressing a specific gene; "pct.2" represents the fraction of cells outside a specific cell type expressing a specific gene.

**Supp. Table S10: Technical summaries of library read processing of the mouse samples**
Technical summaries of computational read processing pipeline for each mouse library.

**Supp. Table S11: Conserved marker table**

Table concatenating the results of the balanced differential testing per cell type including annotation whether a gene was identified as conserved marker in human and mice. The column "species" signifies whether results are derived from a Wilcoxon rank-sum test performed in the human or mouse dataset, the column "dataset" signifies whether the test was performed on the full lung dataset or on the subset of bona fide pulmonary ECs. Each gene is annotated whether it was identified as pan-endothelial, pan-vascular or lymphatic conserved marker gene or a conserved marker gene of the bona fide pulmonary EC subvarieties. If the test was performed on the human dataset, the column "Gene.name.mouse" represents the translated homologue if a one-to-one homologue in the respective other species was available. If there is none one-to-one homologue, it was flagged as "NA" in the "Gene.name.mouse" column. Vice versa, a gene name translation was performed to human genes, when the test was performed on the mouse dataset. "pct.1" represents the fraction of cells within a specific cell type expressing a specific gene; "pct.2" represents the fraction of cells outside a specific cell type expressing a specific gene.
